# Resolving cofactor imbalance in triacylglycerol biosynthesis in oilseeds through glycolytic shunts: a modeling study

**DOI:** 10.1101/2025.07.31.667993

**Authors:** Jorg Schwender

## Abstract

During seed development, carbohydrates are rapidly converted into triacylglycerols (TAGs), with glycolysis and the oxidative pentose phosphate pathway (OPPP) traditionally considered key sources of acetyl-CoA, ATP, NADH, and NADPH for fatty acid synthesis. However, how these classical pathways integrate into an overall stoichiometrically balanced conversion of sugars to TAGs remains unclear. Previous biochemical and isotope-tracing studies in oilseeds across species have revealed that glycolysis is partially bypassed by alternative routes involving ribulose-1,5-bisphosphate carboxylase/oxygenase (RubisCO) and NADPH-dependent malic enzyme (NADPH-ME). The role of these glycolytic shunts in the overall conversion of carbohydrates to TAG is not fully resolved. Here, we evaluate a minimal stoichiometric model for the carbohydrate-to-TAG conversion that satisfies complete cofactor balancing. Conversion solely through glycolysis and OPPP leads to NADH overproduction and cofactor imbalance. Balanced scenarios require inclusion of the malate shunt, the oxidative RubisCO shunt, or an NADPH-producing glyceraldehyde 3-phosphate dehydrogenase variant. All balanced routes also necessitate active mitochondrial oxidative phosphorylation to convert excess NADH to ATP. Applying a large-scale model of central metabolism in developing *Brassica napus* seeds, I further predict that glycolytic bypass usage increases in parallel with seed oil content, supporting their role in enabling high lipid biosynthetic fluxes under physiological constraints.

**Highlight:** Conventional carbohydrate-to-triacylglycerol conversion is cofactor-imbalanced; flux balance analysis shows glycolytic bypasses help correct this, enabling higher biosynthetic rates.

## Introduction

In plants, triacylglycerol (TAG) serves as an energy-dense store of reduced carbon in seeds and other organs, providing the largest source of renewable reduced carbon available from nature (Thelen and Ohlrogge, 2002). TAG consists of three fatty acid molecules bound to a glycerol backbone. In plants, *de novo* fatty acid biosynthesis (FAS) takes place in the plastid compartment from the two-carbon precursor acetyl-CoA by a multi-step process which is highly demanding in energy cofactors. In developing oil seeds, FAS takes place at high rates and sugars provide the main source for biosynthetic precursor and energy cofactors. Acetyl-CoA is provided by sugar catabolic routes (Allen *et al*., 2009; Emes and Neuhaus, 1997; Rawsthorne, 2002; Schwender *et al*., 2004; Schwender and Ohlrogge, 2002). Since FAS and TAG synthesis are highly active in developing oilseeds, the conversion of sugar to TAG is characterized by substantial carbon flux and it is of interest to define and understand this biochemical conversion as an integrated metabolic pathway, in the sense of a well-defined set of reactions that allows conversion of sugar into TAG, while all intermediates and energy cofactors are fully balanced. A clear definition of a minimal set of biochemical reactions that enables a fully balanced conversion should provide a better basis for engineering TAG accumulation in seeds and other tissues.

Although early biochemical studies (Rawsthorne, 2002) and metabolic models (Clark *et al*., 2020) have analyzed the process, a comprehensive, balanced sugar-to-TAG conversion pathway has not been clearly articulated. It is well established that classical glycolysis splits hexose phosphates into pyruvate, and plastidic pyruvate dehydrogenase (PDH) converts pyruvate to acetyl-CoA (Oliver *et al*., 2009). However, a complete conversion scheme that balances acetyl-CoA production along with ATP, NADH, and NADPH requirements is lacking. Conceptually, the transformation can be divided into a catabolic supply side (precursor and cofactor generation) and a biosynthetic demand side (FAS). For simplicity, we consider only plastidial *de novo* FAS, which accounts for most FAS activity in oil-producing tissues (J Browse and Somerville, 1991; Li-Beisson *et al*., 2013; Ohlrogge *et al*., 1979). Regarding the requirement for reducing agents, it is noteworthy that the two sequential [acyl carrier protein] reductase steps require NADH and NADPH as reducing cofactors, respectively (Gonzalez-Thuillier *et al*., 2015; Kater *et al*., 1991; Sheldon *et al*., 1992; Shimakata and Stumpf, 1982a, b; Slabas *et al*., 1986; Vick and Beevers, 1978; Weaire and Kekwick, 1975). Overall, plastidic FAS requires 1 mole of each ATP, NADH and NADPH per mole acetyl-CoA incorporated into a fatty acid chain in the elongation cycle. On the supply side, the classical stoichiometry for glycolysis results in a net gain of 1 mole ATP and 1 mole NADH per mole pyruvate formed (Dennis *et al*., 1997). When pyruvate is next transformed into acetyl-CoA by the plastidic pyruvate dehydrogenase (PDH) (Bao *et al*., 2000; Ke *et al*., 2000; Schwender and Ohlrogge, 2002; Schwender *et al*., 2006), one mole of NADH is added to the one mole of NADH already produced by glycolysis. To cover the NADPH demand, it can be assumed that part of the carbohydrate is oxidized to CO_2_ via the oxidative pentose phosphate pathway (OPPP) (Schuster *et al*., 2000), independent of glycolysis. The OPPP is widely regarded as a principal source of reductant (NADPH) in the absence of photosynthesis for various biosynthetic processes in plants (Kruger and von Schaewen, 2003; Neuhaus and Emes, 2000; Rawsthorne, 2002). While the OPPP-generated NADPH can be used in the fatty acid elongation cycle, one of the two NADH generated along with acetyl-CoA will be in excess. Overall, it seems impossible to find an overall balanced glucose to TAG conversion pathway if we rely solely on glycolysis, PDH, and OPPP on the supply side.

In light of this imbalance, multiple ^13^C-metabolic flux analysis (_13_C-MFA) studies on in vitro–cultured oil-accumulating embryos from *Arabidopsis*, soybean, rapeseed, *Camelina sativa*, sunflower, flax, and maize have provided quantitative insights into central metabolism (Acket *et al*., 2019; Allen *et al*., 2009; Allen and Young, 2013; Alonso *et al*., 2010; Alonso *et al*., 2007; Carey *et al*., 2020; Lonien and Schwender, 2009; Schwender *et al*., 2015; Schwender *et al*., 2003; Sriram *et al*., 2004). Alternative to the OPPP, biochemical and flux studies have shown that plastidic NADP-malic enzyme (NADPH-ME) can contribute both pyruvate and NADPH to FAS in Castor seed endosperm and other oil seeds (Alonso *et al*., 2010; Buchanan *et al*., 2000; Eastmond *et al*., 1997; Neuhaus and Emes, 2000; Rawsthorne, 2002; Smith *et al*., 1992). The proposed pathway bypasses the glycolytic pyruvate kinase step through carboxylation of phosphoenol pyruvate and subsequent reduction of the resulting oxaloacetate to malate by a NADH-dependent malate dehydrogenase. Then, in the plastid, NADPH-malic enzyme delivers pyruvate together with NADPH (Smith *et al*., 1992). Other ^13^C-MFA studies have found that in developing embryos of *Brassica napus* (Schwender *et al*., 2004; Schwender *et al*., 2015) and of other species (Cocuron and Alonso, 2024; Lonien and Schwender, 2009; Tsogtbaatar *et al*., 2020), ribulose-1,5-bisphosphate carboxylase/oxygenase (RubisCO) participates in another partial bypass of glycolytic flux (Schwender *et al*., 2004). Overall, the ^13^C-MFA studies on developing oilseeds in different species suggest that OPPP and malate shunt can cover the NADPH of FAS to varying degrees (O’Grady *et al*., 2012; Sagun *et al*., 2023) and it appears the conversion of sugars to TAG is typically accompanied by significant flux through metabolic bypasses (shunts) of glycolysis, namely via the malate shunt or the RubisCO shunt.

In the light of the problem of NADH excess in the conversion of glucose to TAG based on canonical pathways and given the regular activity of glycolytic shunts found in the empirical ^13^C-MFA studies, the question arises whether these extensions of the canonical glycolytic and OPPP routes are simply auxiliary or essential components of a balanced sugar-to-TAG pathway. To explore this question, one could use large-scale stoichiometric models such as *Bna572+*, a metabolic stoichiometric model with more than 600 reactions representing seed primary metabolism for *Brassica napus* developing seeds (Hay *et al*., 2014; Hay and Schwender, 2011b; Schwender and Hay, 2012). All of the just mentioned relevant biochemical reactions are present in *Bna572+*. However, due to the complexity of the model with the presence of subcellular compartments with duplication of glycolysis and other pathways in the cytosol and plastid, the enumeration of all possible conversions of sugar to TAG leads to an unmanageable number of independent pathways that must be analyzed. For this reason, here I present a small-scale uncompartmentalized model containing glycolysis, PDH, OPPP as well as the relevant glycolytic shunts and analyze fully balanced pathways of glucose to TAG conversion. First, it is clearly demonstrated that, for a minimal configuration where only glycolysis, PDH and OPPP are active in producing acetyl-CoA, glycerol 3-phosphate and energy cofactors, the overall conversion results in a substantial NADH surplus arising from glycolysis. The activity of the malate shunt, the oxidative RubisCO shunt or of an NADPH dependent isoform of the glycolytic enzyme glyceraldehyde 3-phosphate dehydrogenase each enable balanced conversion schemes. In each case the NADH surplus is avoided in essence by bypassing the NADH-dependent glyceraldehyde 3-phosphate dehydrogenase step of glycolysis or by transhydrogenation of the glycolytic NADH onto NADP (malate shunt). Furthermore, in all three balanced schemes mitochondrial oxidative phosphorylation is required to contribute ATP. The activity of the three glycolytic shunts is then also examined for simulations of *Bna572+*, an FBA model for central metabolism in developing *B. napus* embryos.

## Materials and methods

### Constraint-based modeling

Flux modeling was performed using the constraint-based reconstruction and analysis (COBRA) toolbox version 3.1 (Heirendt *et al*., 2019) and GLPK solver (https://www.gnu.org/software/glpk/) within the MATLAB R2021a environment (The MathWorks, Natick, MA, United States). The basic simulation procedure used here is to use Flux Balance Analysis (FBA) (Orth *et al*., 2010) to predict cellular flux states that satisfy an optimality criterion. Bilevel optimization is applied by first solving the overall metabolic state by minimizing substrate uptakes relative to a fixed biomass accumulation rate and applying this optimum as a new constraint to the model. Then the inner flux solution space is further explored by maximizing and minimizing the rate of inner reactions (Flux Variability Analysis). To detect cofactor imbalances, ATP surplus was quantified by maximizing an ATP-consuming dummy reaction in a secondary optimization (Clark and Schwender, 2022). Another case of secondary optimization was applied to detect the flux solution space for glycolytic shunts, which are defined as combinations of reactions already present in the model. To do this, the summary stoichiometry of network reactions that represents the glycolytic shunt was added as a new reaction to the network and then minimized or maximized.

In addition to the analysis with the COBRA toolbox, the extreme pathway software tool X3 (Thiele and Palsson, 2010) was used to enumerate independent conversion pathways. The software tool was downloaded from https://systemsbiology.ucsd.edu/Downloads/ExtremePathwayAnalysis and executed from the DOS command line.

### Simulation of small-scale stoichiometric model

Possible conversion pathways of glucose into TAG were predicted by extreme pathway analysis (Papin *et al*., 2003; Schilling *et al*., 2000). As in FBA, the steady-state solution space is limited by the reaction stoichiometries and thermodynamic irreversibility constraints. Extreme pathway analysis enumerates all independent pathways (extreme pathways) that constitute the edges of the convex cone of the solution space. All possible flux distributions in a model can be represented as linear combinations of extreme pathways. Note that, by definition, extreme pathways are non-decomposable (Papin *et al*., 2003) which means that an extreme pathway cannot remain operational if any of the active reactions are removed. To calculate extreme pathways, a given COBRA model structure was translated into the input file for the extreme pathway software tool X3 using the COBRA function *convertModelToEX*. In addition to this exploration of independent pathways, specific extreme pathways were re-calculated by a two-level optimization using the COBRA function *optimizeCbModel*. First, the glucose uptake flux (reaction Glc_up) was minimized for a fixed rate of TAG synthesis. After fixing the optimized uptake rate in the model a secondary reaction that is characteristically active in the extreme pathway was maximized.

### Carbon Conversion Efficiency

For a given flux solution, the total amount of carbon uptakes and carbon leaving the system (biomass) is balanced. Typically, CO_2_ efflux is the only way carbon leaves the system, besides biomass. Therefore, the difference between the total carbon uptake and the carbon efflux as CO_2_, divided by the total carbon uptake equals the fraction of carbon that was converted into biomass (Carbon Conversion Efficiency, CCE).

### Chemical limit for the conversion of glucose into TAG

Independent of specific metabolic pathways, for any biochemical process converting glucose into a carbon reduced product a chemical conversion stoichiometry can be derived based on the chemical and redox balance. The conversion efficiency of the chemical process defines the best possible conversion by any given metabolic pathway (chemical limit). A general calculation scheme for the chemical limit of a conversion of glucose into TAG is described in supplemental Material 1.

### Variability in respiratory ATP yield in the small-scale model

In the small-scale model, mitochondrial oxidative phosphorylation is described by a summary stoichiometry (reaction *mitoOP*). To model different values for the ratio of ATP produced per NADH consumed by oxidative phosphorylation (P/O ratio) the coefficients of the reaction mitoOP were modified by the multiplier “PO” in the following way:

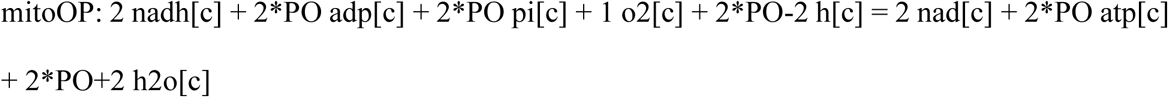

### Modeling of oilseed metabolism by Flux Balance Analysis with a large-scale model

To simulate oilseed metabolism *Bna572+* was used, a large scale metabolic stoichiometric model with 669 reactions representing seed primary metabolism for *Brassica napus* developing seeds (Hay *et al*., 2014; Hay and Schwender, 2011b; Schwender and Hay, 2012). *Bna572+* in COBRA-compatible Systems Biology Markup Language (SBML) can be accessed online (https://doi.org/10.3389/fpls.2014.00724, Supplemental material, DataSheet1.zip, file name “bna20140218T082916.xml”). Cellular flux states were predicted by FBA by minimization of the total carbon molar substrate uptake (objective function) relative to a fixed biomass flux. The assumption that metabolism is overall carbon-efficient is supported by the physiology of the developing seed. Given that in most Brassicaceae the embryo developing inside a seed receives reduced carbon substrates from its endosperm environment and that seed coat and silique walls are likely a strong diffusion barrier for respired CO_2_ (Goffman *et al*., 2005; Goffman *et al*., 2004), one can assume that losses of carbon as respiratory CO_2_ are kept to a minimum and that the highest possible conversion efficiency is maintained.

### Adjustments of irreversibility constraints in Bna572+

In previous model studies of seed metabolism using FBA with *Bna572+*, the oxidative pentose phosphate pathway (OPPP) was typically predicted to be inactive (Hay and Schwender, 2011a; Schwender and Hay, 2012). This outcome appears unrealistic, given that parallel experimental studies employing ¹³C-metabolic flux analysis in developing *Brassica napus* embryos consistently revealed substantial OPPP flux (Hay *et al*., 2014; Schwender *et al*., 2015; Schwender *et al*., 2003). The failure to predict OPPP activity has also been noted in other plant FBA models. Notably, it has been demonstrated that OPPP flux can be recovered by eliminating transhydrogenase cycles that permit unrestricted transfer of reducing equivalents from NADH to NADP⁺ (Cheung *et al*., 2013; Sweetlove *et al*., 2013). Such transfers, when uncoupled from a driving thermodynamic force, are considered biochemically implausible. As a result, these transhydrogenation schemes can be blocked in FBA models by adjusting irreversibility constraints (Cheung *et al*., 2013). Prompted by these insights, the network topology of *Bna572+* was re-examined. Two distinct NADH-to-NADP⁺ transhydrogenase routes were identified (Fig. S1). It was shown that preventing the operation of these energetically neutral transhydrogenase cycles enables the model to predict OPPP flux—provided that the following two specific reactions are constrained: phosphoenolpyruvate carboxykinase (PPCKc; EC 4.1.1.32) and mitochondrial NADH-dependent isocitrate dehydrogenase (IDHm). These enzymes must be restricted from operating in the direction of phosphoenolpyruvate (PEP) carboxylation and 2-oxoglutarate (α-ketoglutarate) reduction, respectively (Table S1). Introducing these two constraints is consistent with established physiological knowledge. For PPCK, in all documented cases where the enzyme plays a functional role—such as during photosynthesis, C₄ and Crassulacean acid metabolism, and gluconeogenesis in germinating oilseeds—it catalyzes the decarboxylation of oxaloacetate to form PEP (Rojas and Iglesias, 2023). Although in vitro studies have confirmed full reversibility of the reaction, the biochemical characteristics of the plant enzyme support a predominantly decarboxylating direction in vivo (Rojas *et al*., 2021). Therefore, modifying the model to enforce unidirectionality in the decarboxylation direction for PPCK is appropriate. Regarding isocitrate dehydrogenase, the enzyme is only required to function in the oxidative direction (isocitrate to 2-oxoglutarate) for the tricarboxylic acid (TCA) cycle to proceed. However, prior ¹³C-glutamine labeling experiments in *B. napus* embryos have shown that isocitrate dehydrogenase can be reversible *in vivo* (Schwender *et al*., 2006). Consequently, all isoforms of isocitrate dehydrogenase in *Bna572+* were retained as bidirectional. To block transhydrogenase cycling specifically, only the mitochondrial NADH-dependent isoform (IDHm) was constrained to operate exclusively in the oxidative (forward) direction. This restriction is justified by the biochemical properties of mitochondrial NADH-dependent IDH, which indicate that the enzyme functions solely in the forward direction (producing NADH) under physiological conditions (Igamberdiev and Gardeström, 2003).

### Model reduction to identify glucose-to-TAG conversion pathways (bna572+)

To explore different pathways for the conversion of glucose to TAG, the FBA model *Bna572+*, which represents the metabolism in a developing *Brassica napus* oilseed, was used as a starting point. The model was configured as previously described (Hay *et al*., 2014), except that reactions PPCKc and IDHm were set to be irreversible (positive rates permitted) to disable no-cost NADH-to-NADP⁺ transhydrogenation. To adapt the network to simulate only biosynthesis of tristearoylglycerol from glucose, most reactions enabling exchange with the external environment were deactivated, permitting only glucose uptake, oxygen/CO₂ exchange and TAG export. The biomass synthesis was replaced by a reaction equation that summarizes the biosynthesis of stearoyl-CoA (reaction FAS) and one summary stoichiometry for TAG biosynthesis (Reaction TAGSynth). Since maintenance requirements do not contribute to biosynthesis, the maintenance ATP demand reaction (ATPdrain_c) was disabled. Next flux solutions were calculated for four different optimization criteria: Maximize TAG production, minimize oxygen uptake, minimize total ATP production and minimize glucose uptake. In all four cases the obtained flux solution was characterized by same glucose uptake rate and CCE, showing that the conversion efficiency is not depending on one specific optimization criterion. With the optimum glucose uptake (Ex_glc_a) fixed relative to the TAG production rate, the model was reduced by removing inactive reactions from the model based on the flux solution space determined by Flux Valiability Analysis. The reduced model is summarized in Table S2.

### Physiological constraints for prediction of whole seed metabolism with bna572+

*Bna572+* was configured to simulate photoheterotrophic growth conditions of *in vitro* cultured embryos from *Brassica napus* genotype 3170, as previously described by Hay *et al*. (Hay *et al*., 2014). That earlier study introduced the *Bna572+* FBA model alongside a comprehensive ¹³C-metabolic flux analysis (¹³C-MFA) of cultured *B. napus* embryos. The physiological constraints used in the current study were based on those original parameters, with minor adjustments (Table 1). Whereas prior simulations accounted for cellular maintenance solely through ATP consumption via a generic ATPase reaction, the current model also incorporates redox (NADPH) maintenance requirements. This addition follows findings by Cheung *et al*. (2013), who demonstrated that attributing maintenance energy demands to both ATP and NADPH consumption improved alignment between FBA predictions and empirical flux measurements in plant cell cultures (Cheung *et al*., 2013). To implement this, a generic NADPH oxidation reaction (*NADPHdrain_c*) was added to the network, and the model was adjusted using a procedure adapted from Cheung et al. In brief, Cheung et al. simulated a tradeoff between ATP and NADPH maintenance costs by constructing a Pareto front—a curve that explores various combinations of ATP and NADPH consumption that match the experimentally determined glucose uptake rate. While maintaining constant carbon conversion efficiency along this front, they observed variations in internal flux distributions. By identifying the point where both glucose uptake and the ratio of OPPP flux to glycolysis flux aligned with ¹³C-MFA results, they determined optimal maintenance cost values. This same principle was applied here to *Bna572+*, with an important modification: due to the photoheterotrophic nature of the cultured embryos, the impact of ATP and NADPH generation from photosynthetic light reactions was also considered in the cofactor balance. The resulting Pareto front for a representative light flux of 5 µmol photons/h is shown in Fig. S2. At this light input, the model best matches experimental data with an ATP maintenance flux of 1.82 µmol/h and an NADPH maintenance flux of 0.42 µmol/h, reproducing a carbon conversion efficiency of 81.1% and a relative OPPP flux (OPPP flux/total hexose uptake flux) of 0.16, in line with the ¹³C-MFA observations (Table 1). Note that the photosynthetic light flux and associated ATP and NADPH maintenance costs cannot be directly measured. However, Fig. S3A demonstrates that the same agreement with the ¹³C-MFA data can be achieved across a range of assumed light flux levels. Essentially, the model can be tuned to match experimental results by balancing photosynthetic cofactor production with appropriate maintenance demands. Specifically, the ATP maintenance requirement must exceed the photosynthetic ATP supply by 0.213 µmol/h, while the NADPH maintenance flux must fall 0.833 µmol/h short of photosynthetic NADPH production (Fig. S3A). As shown in Fig. S3B, depending on light availability, ATP maintenance costs account for 29–47% of total ATP production, whereas NADPH maintenance demands comprise 0–37% of total NADPH output.

**Table 1.**
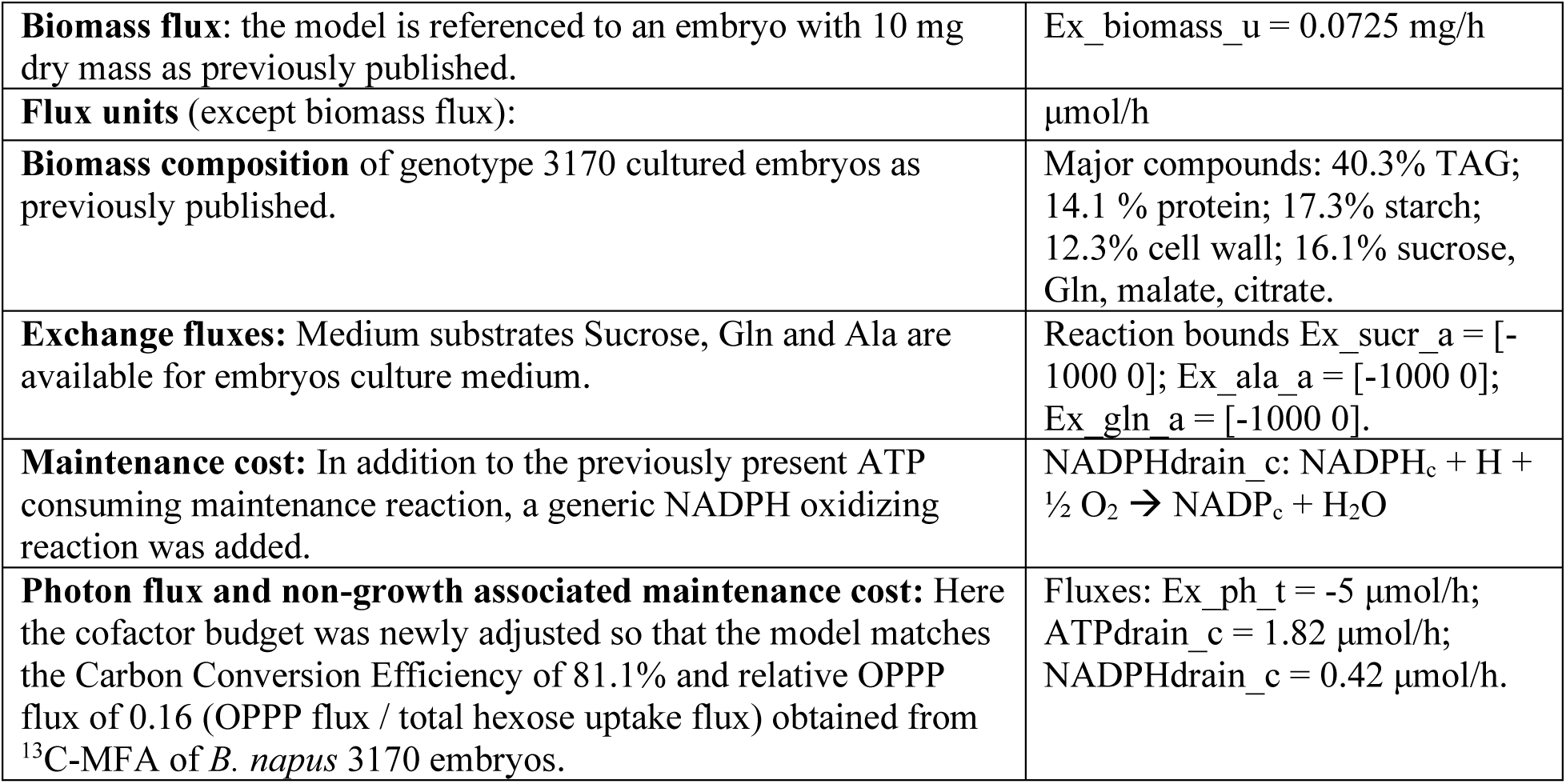
Adjustments of *Bna572+* to represent *Brassica napus* cultured embryos (genotype 3170). The model was based on original parameters as published previously (Hay *et al*., 2014) with minor adjustments.

|  |  |
| --- | --- |
| **Biomass flux:** the model is referenced to an embryo with 10 mg dry mass as previously published. | Ex_biomass_u = 0.0725 mg/h |
| **Flux units** (except biomass flux): | μmol/h |
| **Biomass composition** of genotype 3170 cultured embryos as previously published. | Major compounds: 40.3% TAG; 14.1 % protein; 17.3% starch; 12.3% cell wall; 16.1% sucrose, Gln, malate, citrate. |
| **Exchange fluxes:** Medium substrates Sucrose, Gln and Ala are available for embryos culture medium. | Reaction bounds Ex_sucr_a = [-1000 0]; Ex_ala_a = [-1000 0]; Ex_gln_a = [-1000 0]. |
| **Maintenance cost:** In addition to the previously present ATP consuming maintenance reaction, a generic NADPH oxidizing reaction was added. | NADPHdrain_c: $\text{NADPH}_c + \text{H} + \frac{1}{2} \text{O}_2 \rightarrow \text{NADP}_c + \text{H}_2\text{O}$ |
| **Photon flux and non-growth associated maintenance cost:** Here the cofactor budget was newly adjusted so that the model matches the Carbon Conversion Efficiency of 81.1% and relative OPPP flux of 0.16 (OPPP flux / total hexose uptake flux) obtained from <sup>13</sup>C-MFA of *B. napus* 3170 embryos. | Fluxes: Ex_ph_t = -5 μmol/h; ATPdrain_c = 1.82 μmol/h; NADPHdrain_c = 0.42 μmol/h. |

### Test for presence of no-cost transhydrogenase schemes

To assess whether a FBA model permits cost-free transhydrogenation—i.e., the transfer of reducing equivalents from NADH to NADP⁺ without incurring a net ATP cost— the following method was devised. Two generic reactions were introduced into the model: (1) *GnTH*, representing direct transfer of reducing power between the NAD(H) and NADP(H) pools, and (2) *GnATPsynth*, representing ATP resynthesis from ADP and inorganic phosphate. I then conducted a series of sensitivity analyses, in which the objective function *Z* was evaluated in response to small perturbations in the fluxes through *GnTH* and *GnATPsynth*. Specifically, small changes ∂NADPH and ∂ATP corresponding to a small amount of NADPH formation via transhydrogenation and ATP synthesis were indroduced, respectively. The sensitivities of the objective function to these changes were defined as:

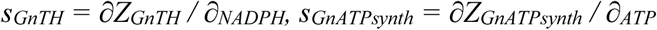

Of particular interest is then the condition under which ATP resynthesis energetically compensates for NADH-to-NADPH transhydrogenation. This occurs when the changes in the objective function are equal in magnitude but opposite in sign:

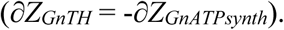

Substituting the sensitivities yields:

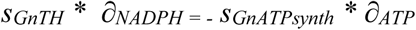

Rearranging gives the ATP cost per mole of NADPH transhydrogenated:

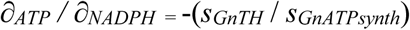

This ratio quantifies the energetic cost, in terms of ATP, associated with NADPH generation via transhydrogenation within the modeled metabolic network.

### Model availability

The small-scale model for conversion of glucose to TAG and the seed model *Bna572+* are provided with simulation codes in Supplementary File 2.

## Results and discussion

Following the experimental findings on OPPP flux and glycolytic shunt activities made in ¹³C-MFA studies on oilseed development, it is of interest to reproduce these results within a predictive metabolic modeling framework. However, when employing *Bna572+* - a large-scale reconstruction of primary seed metabolism in *Brassica napus* developing seeds (Hay *et al*., 2014; Hay and Schwender, 2011b; Schwender and Hay, 2012) - the resulting flux predictions diverged from those inferred via ¹³C-MFA, particularly regarding the failure of FBA to predict oxidative pentose phosphate pathway (OPPP) activity (Hay *et al*., 2014; Schwender *et al*., 2015; Schwender *et al*., 2003). As detailed in the “Materials and Methods” section, this discrepancy can now be attributed to the network architecture of *Bna572+*, which permitted the transfer of reducing equivalents from NADH to NADP⁺. This mechanism effectively bypassed the direct generation of NADPH via oxidative decarboxylation in the OPPP. By imposing stricter irreversibility constraints on select reactions, such NADH-to-NADP⁺ transhydrogenation was disabled, thereby allowing the model to predict flux through the OPPP for NADPH production (see “Materials and Methods”). With this refined FBA model, I investigated the metabolic potential for converting glucose into triacylglycerol (TAG) in greater detail. The *Bna572+* network was adapted to simulate TAG biosynthesis from glucose, excluding the synthesis of other biomass constituents or byproducts. To this end, most reactions enabling exchange with the external environment were deactivated, permitting only glucose uptake and oxygen/CO₂ exchange. The model was constrained to produce only tristearoylglycerol, and the maintenance ATP demand (ATPdrain_c) was disabled. Subsequently, the model was reduced by eliminating reactions that carried zero flux under these conditions, resulting in a simplified network of 91 active reactions (from the original 671, Table S2). The reduced network represents all possible ways the network can carry out the overall conversion of glucose to TAG efficiently. Key active pathways of intermediary metabolism included glycolysis, pyruvate dehydrogenase (PDH), the OPPP, NADP-malate dehydrogenase, phosphoribulokinase, and RubisCO (Table S2). Within mitochondrial metabolism, oxidative phosphorylation remained functional, whereas the tricarboxylic acid (TCA) cycle was inactive. Notably, the non-phosphorylating NADPH-dependent glyceraldehyde-3-phosphate dehydrogenase (ALDH_c) was also active in the model, suggesting contribution of this enzyme to TAG synthesis. To further delineate distinct pathways of glucose-to-TAG conversion, extreme pathway analysis was applied to the 91-reaction network. This resulted in well over 200,000 predicted pathways and it became clear that the compartmentation of the model and parallelism of reactions in cytosol and plastid were largely responsible for this complexity. Therefore, it seemed sensible to cast the network of reactions that are active in the conversion of glucose into TAG into an even simpler version.

### Defining a small-scale model for the conversion of sugars to TAG in plants

A small-scale primary metabolism network was defined based on the 91-reaction network (Table S2) while removing most of its complexity due to compartmentalization. The model (Fig. 1) can convert glucose into tristearate, CO_2_ and H_2_O, while oxygen can also enter to be consumed by respiratory functions. Relative to the 91-reaction network, the canonical glycolysis with its ATP dependent kinase reactions (Plaxton, 1996) were retained while pyrophosphate dependent glycolytic reactions (reactions PFP_c, PPDK_c) were omitted. To generate NADPH, a complete set of OPPP reactions is present to enable the pathway in both non-cyclic and cyclic mode (Schuster *et al*., 2000) (Fig. 1). In addition, the malate shunt (Smith *et al*., 1992), which bypasses the glycolytic pyruvate kinase (PK) step and generates pyruvate via NADPH-malic enzyme, is also included (Fig. 1, green). Also included was the non-phosphorylating NADPH-dependent glyceraldehyde-3-phosphate dehydrogenase that was active in the 91-reaction network, abbreviated in the small-scale network as “npGAPDH” (Fig. 1). Besides the alternatives for NADPH generation, adding phosphoribulokinase (PRK) and RubisCO to the network allows operation of the oxidative and non-oxidative modes of the RubisCO shunt (Schwender *et al*., 2004) (Fig. 1). The process of oxidative phosphorylation in mitochondria was summarized in one reaction (*mitoOP*, Fig. 1), with the efficiency conservatively estimated as an integer of 2 ATP/NADH (P/O ratio) (Lambers, 1997). To capture stoichiometries with excess generation of ATP, a generic ATP hydrolyzing reaction (*ATPsurplus*, Fig. 1) was included in the small-scale model. Scenarios that include ATPsurplus flux are thought to link to an external activity that hydrolyzes ATP so that the total process can continue. Without that external process the conversion cannot be balanced. On the demand side, FAS is included in the model as the summary stoichiometry for the synthesis of stearic acid as derived before (Clark and Schwender, 2022). The FAS summary stoichiometry accounts for the plastidic *de-novo* synthesis of stearic acid and integrates the ATP cost for activation of the free fatty acid to the CoA ester by long-chain acyl-CoA synthetase (EC 6.2.1.3) in the cytosol/ER (Shockey *et al*., 2002). The resulting CoA ester can be incorporated into glycolipids without additional energy expense (Li-Beisson *et al*., 2013) and TAG assembly is simplified into a lumped reaction (Fig. 1).

**Fig. 1:**
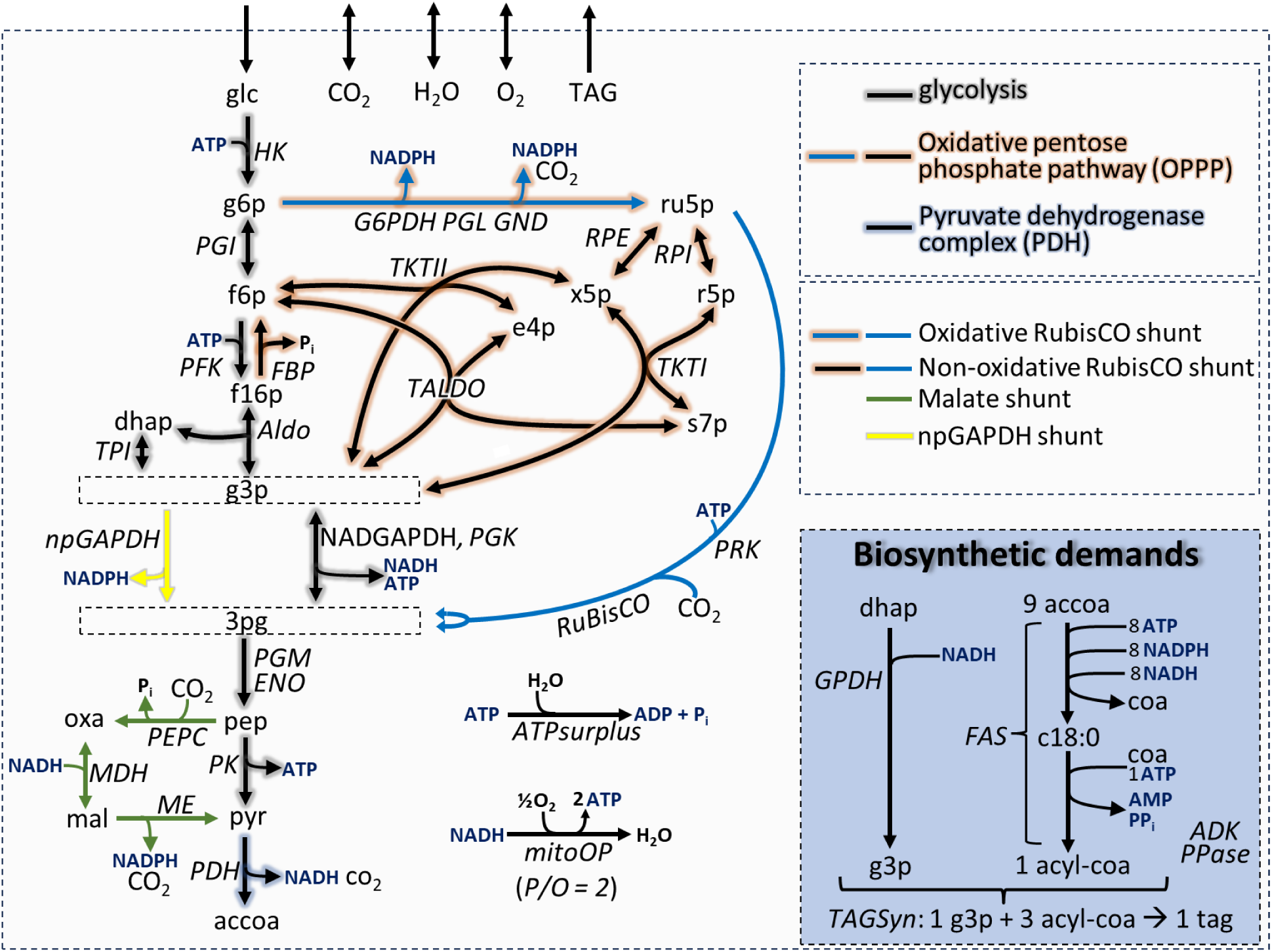
Schematic of a small-scale stoichiometric network representing the conversion of glucose (glc) to triacylglycerol (TAG) in plants. Names of reactions are shown in italics. Detailed reaction stoichiometries are listed in Table S3 and metabolites are identified in Table S4. Enzymes: ADK, Adenylate kinase EC 2.7.4.3; Aldo, Fructose bisphosphate aldolase EC 4.1.2.13; ATPsurplus: generic ATP hydrolyzing reaction; Eno, enolase EC 4.2.1.11; FAS, combined reaction stoichiometries to generate C18:0-CoA from acetyl-CoA and cofactors; Fbp, Fructose bisphosphatase; EC 3.1.3.11; GPDH, glycerol-3-phosphate dehydrogenase (NAD+); EC 1.1.1.8; G6PDH, glucose-6-phosphate dehydrogenase EC 1.1.1.49; GND, 6-phosphogluconate dehydrogenase EC 1.1.1.44; HK, hexokinase EC 2.7.1.2; MDH, malate dehydrogenase EC 1.1.1.37; ME, malic enzyme EC 1.1.1.40; mitoOP, oxidative phosphorylation, 2 ATP per NADH; NADGAPDH, NADP-GAP-dehydrogenase EC 1.2.1.13; npGAPDH, NADP-GAP-dehydrogenase, non-phosphorylating EC 1.2.1.9; PDH, pyruvate dehydrogenase EC 1.7.1.4/EC 2.3.1.12/EC 1.2.4.1; PEPC, pep carboxylase EC 4.1.1.31 and carbonic anhydrase EC 4.2.1.1; PFK, Phosphofructokinase EC 2.7.1.11; PGI, Glucose-6P isomerase EC 5.3.1.9; PGK, phosphoglycerate kinase EC 2.7.2.3; PGL, 6-phosphoglucono-lactonase EC 3.1.1.31; PGM, phosphoglycerate mutase EC 5.4.2.11; PK, pyruvate kinase EC 2.7.1.40; PPase, Inorganic pyrophosphatase EC 3.6.1.1; PRK, Phosphoribulokinase EC 2.7.1.19; Riso, ribose-5P isomerase EC 5.3.1.6; RuBisCO, Ribulose bisphosphate carboxylase/oxygenase EC 4.1.1.39; TA, transaldolase EC 2.2.1.2; TKI, transketolase EC 2.2.1.1; TKII, transketolase EC 2.2.1.1; TPI, Triosephosphate isomerase EC 5.3.1.1; Xepi, xylulose-5-phosphate epimerase EC 5.1.3.1.

Altogether, the small-scale model (Fig. 1) consists of 33 reactions and 39 metabolites of which three can be freely exchanged with the environment, glucose can only be taken up and TAG can only be released (see details in Tables S1, S2). To explore all possible balanced conversion pathways of glucose to TAG, extreme pathway analysis was used (Schilling *et al*., 2000), predicting 5 distinct routes for the conversion of glucose into TAG (Table 2, Table S5). The pathways differ in their overall carbon conversion efficiency, in their use of different dehydrogenases to produce NADPH for FAS and the presence or absence of an ATP surplus (Table 2).

**Table 2:**
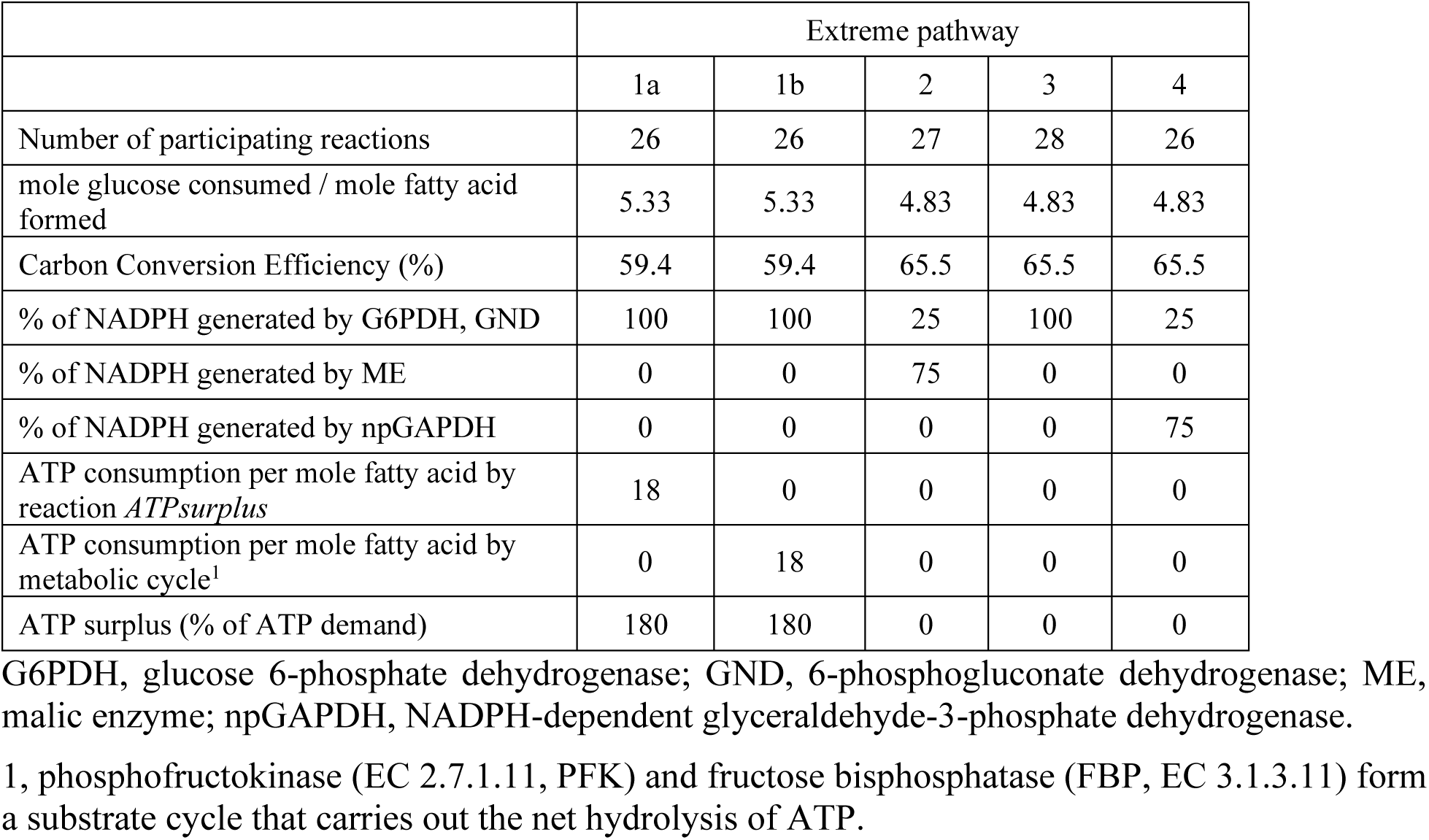
Summary of extreme pathways that describe the conversion of glucose to tristearate in the small-scale model of Fig. 1. The detailed flux results can be found in Supplemental Table S5.

|  | Extreme pathway |  |  |  |  |
| --- | --- | --- | --- | --- | --- |
|  | 1a | 1b | 2 | 3 | 4 |
| Number of participating reactions | 26 | 26 | 27 | 28 | 26 |
| mole glucose consumed / mole fatty acid formed | 5.33 | 5.33 | 4.83 | 4.83 | 4.83 |
| Carbon Conversion Efficiency (%) | 59.4 | 59.4 | 65.5 | 65.5 | 65.5 |
| % of NADPH generated by G6PDH, GND | 100 | 100 | 25 | 100 | 25 |
| % of NADPH generated by ME | 0 | 0 | 75 | 0 | 0 |
| % of NADPH generated by npGAPDH | 0 | 0 | 0 | 0 | 75 |
| ATP consumption per mole fatty acid by reaction *ATPsurplus* | 18 | 0 | 0 | 0 | 0 |
| ATP consumption per mole fatty acid by metabolic cycle<sup>1</sup> | 0 | 18 | 0 | 0 | 0 |
| ATP surplus (% of ATP demand) | 180 | 180 | 0 | 0 | 0 |

G6PDH, glucose 6-phosphate dehydrogenase; GND, 6-phosphogluconate dehydrogenase; ME, malic enzyme; npGAPDH, NADPH-dependent glyceraldehyde-3-phosphate dehydrogenase.
1, phosphofructokinase (EC 2.7.1.11, PFK) and fructose biphosphatase (FBP, EC 3.1.3.11) form a substrate cycle that carries out the net hydrolysis of ATP.

### Glycolysis-based conversion of glucose to fatty acids in plants is inherently imbalanced

In pathways 1a and b (Table 2), glucose is converted into TAG with a Carbon Conversion Efficiency (CCE) of 59.4 %, which is below the chemical limit for the formation of tristearate from glucose of 69.9% (supplemental Material 1). As an explanation for this low efficiency, in both pathways 1a and b an ATP surplus is present that amounts to 180% of the biosynthetic demand (Table 2). In the broader context of cellular metabolism, this ATP surplus could benefit other processes. If the ATP surplus is not consumed by another process, the conversion would stall. In this sense, the overall transformation of pathways 1a and 1b can be considered unbalanced. The same imbalance is also contained in pathway 1b (Table 2), in which case the same net hydrolysis of ATP is caused by a substrate cycle by fructose bisphosphatase (FBP, EC 3.1.3.11) and phosphofructokinase (EC 2.7.1.11, PFK). To provide more insight into the cofactor imbalance, the flux distribution of pathway 1a is summarized in Fig. 2A, with fluxes normalized to 1 mole fatty acid or 1/3 mole TAG. Fig. 2A shows that in the supply section 18 moles NADH are produced along with moles 9 acetyl-CoA. However, only 8.33 moles NADH are needed in the biosynthesis of TAG (Fig. 2A). The remainder is consumed by oxidative phosphorylation (reaction *mitoOP*) and the balance between ATP investment in glycolysis, biosynthetic ATP demand, and the ATP production by glycolysis and oxidative phosphorylation results in the surplus of 18 ATP (Fig. 2A). This means that the cause for the ATP surplus is overproduction of glycolytic NADH that is discharged by oxidative phosphorylation. Again, in the broader context of seed metabolism, this surplus could be consumed by other biosynthetic activities that take place at the same time, such as storage protein synthesis. However, if this were the case, the conversion of sugar to oil would be tightly coupled with this other process and therefore could not easily be controlled independently. In other words, whatever the other ATP consuming process is, it had to be included in the definition of a glucose to TAG conversion pathway.

**Fig. 2:**
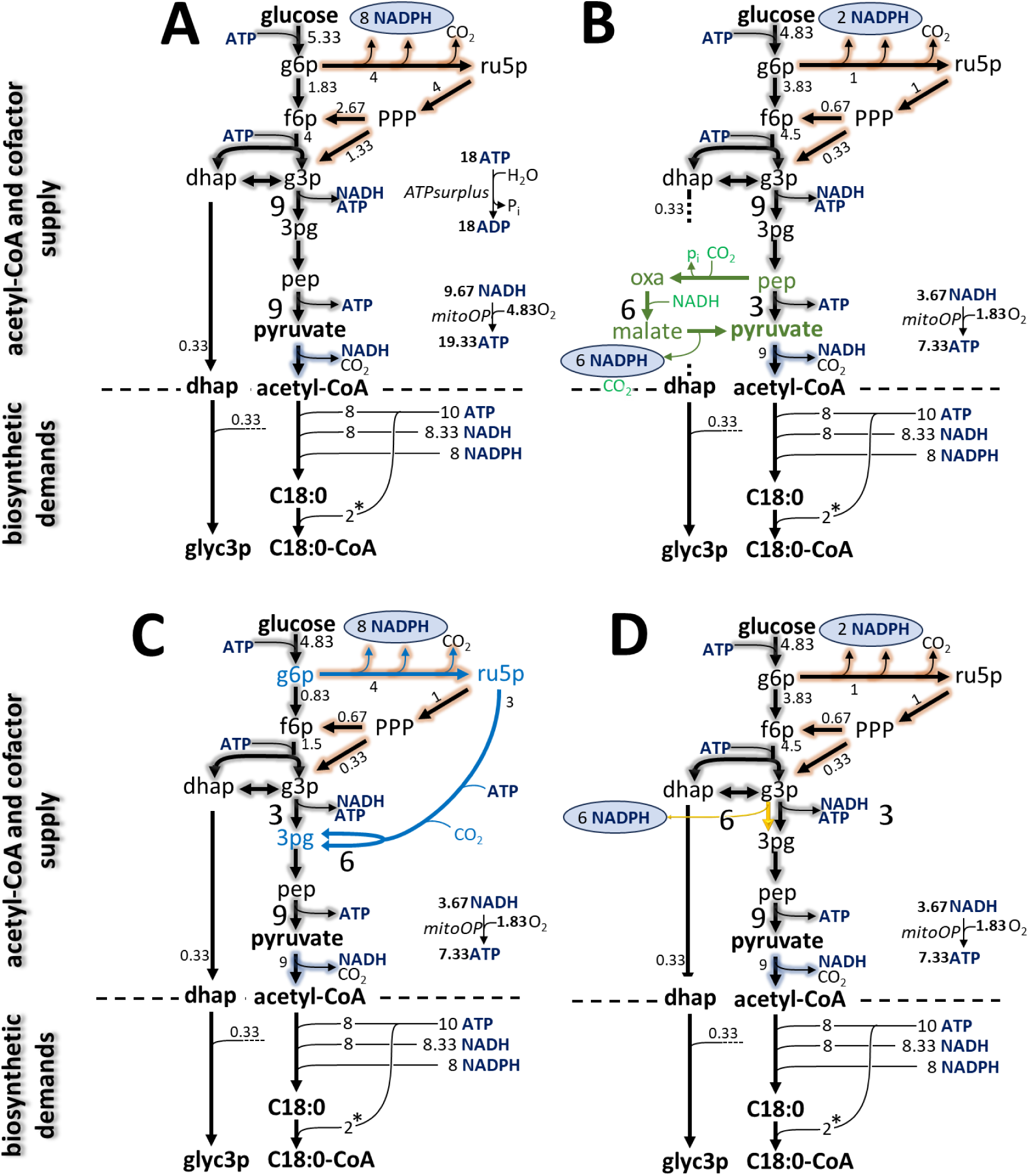
Conversion pathways from glucose to tristearate. Scenarios are predicted by extreme pathway analysis of the small-scale network (Fig. 1). In each panel, only active reactions are shown. Flux units are relative to the formation of one mole stearoyl-CoA (C18:0) and ⅓ mole glycerol 3-phosphate (glyc3p), which combine to ⅓ mole TAG. (**A**), if only the core reactions of glycolysis, pyruvate dehydrogenase and oxidative pentose phosphate pathway are active, a severe ATP surplus results. The ATP surplus is avoided by activity of the malate shunt with transhydrogenase activity (**B**), the oxidative RubisCO shunt (**C**), or non-phosphorylating glyceraldehyde 3-phosphate dehydrogenase (**D**). Large flux numbers highlight proportions of flux meeting at the pyruvate node (**B**) or the 3pg node (**C**, **D**), respectively. The proportions 6:3 mean that 66.7 of glycolytic flux is bypassed. *The cost of acylation of C18:0 to C18:0-CoA by long-chain acyl-CoA synthetase (EC 6.2.1.3) in the cytosol/ER is included as 2 ATP/C18:0. Abbreviations: dhap, dihydroxyacetone phosphate; 3pg, 3-phosphoglycerate; f6p, fructose 6-phosphate; g3p, glyceraldehyde 3-phosphate; g6p, glucose 6-phosphate; mitoOP, mitochondrial oxidative phosphorylation; oxa, oxaloacetate; pep, phosphoenolpyruvate; p_i_, inorganic phosphate; PPP, pentose phosphate pathway; ru5p, ribulose 5-phosphate. Detailed flux results can be found in Supplemental Table S5.

### Glycolytic shunts enable NADH-balanced conversion of glucose to TAG

In contrast to pathways 1a and 1b, pathways 2 to 4 are balanced and have higher conversion efficiency (Table 2). In addition to the core activities of glycolysis, PDH and OPPP, pathways 2 to 4 utilize metabolic shunts of glycolysis that are closer examined in Fig. 2 B-D. In pathway 2, 2/3 of the flux between 3pg and acetyl-CoA bypasses the glycolytic pyruvate kinase step via the malate shunt (Fig. 2B in green). Compared to the unbalanced scenario, NADPH production in the OPPP is reduced and instead 6 mol of NADPH is produced by transhydrogenation of glycolytic NADH to NADP in the malate shunt (Fig. 2B). Compared to the conversion of pep to pyruvate by pyruvate kinase, the malate shunt does not generate ATP (Fig. 2B), so it can be said that the malate shunt has an additional driving force for transhydrogenation.

In pathway 3 (Fig. 2C), part of glycolysis is bypassed by the oxidative RubisCO shunt, which is highlighted in blue as the conversion of glucose-6-phosphate (g6p) to 3-phosphoglycerate (3pg) via ribulose-5-phosphate (ru5p) (Fig. 2C). By re-fixing CO_2_ that was generated by oxidative decarboxylation of g6p, the shunt achieves the same amount of 2 moles 3pg per mole 6pg as the parallel glycolysis section. However, via the oxidative RubisCO shunt, NADH production by NADH-GAPDH is bypassed, thus avoiding the production of 6 mol of NADH and replacing it with 6 mol of NADPH made by oxidation of g6p. While the oxidative RubisCO shunt of glycolysis was formerly described alongside the non-oxidative RubisCO shunt (Schwender *et al*., 2004). Both variants are compared in Fig. 3. The non-oxidative RubisCO shunt is known to be relevant in a photomixotrophic setting (Schwender *et al*., 2004) and is not active in the extreme pathways predicted based on a heterotrophic context (Fig. 2).

**Fig. 3.**
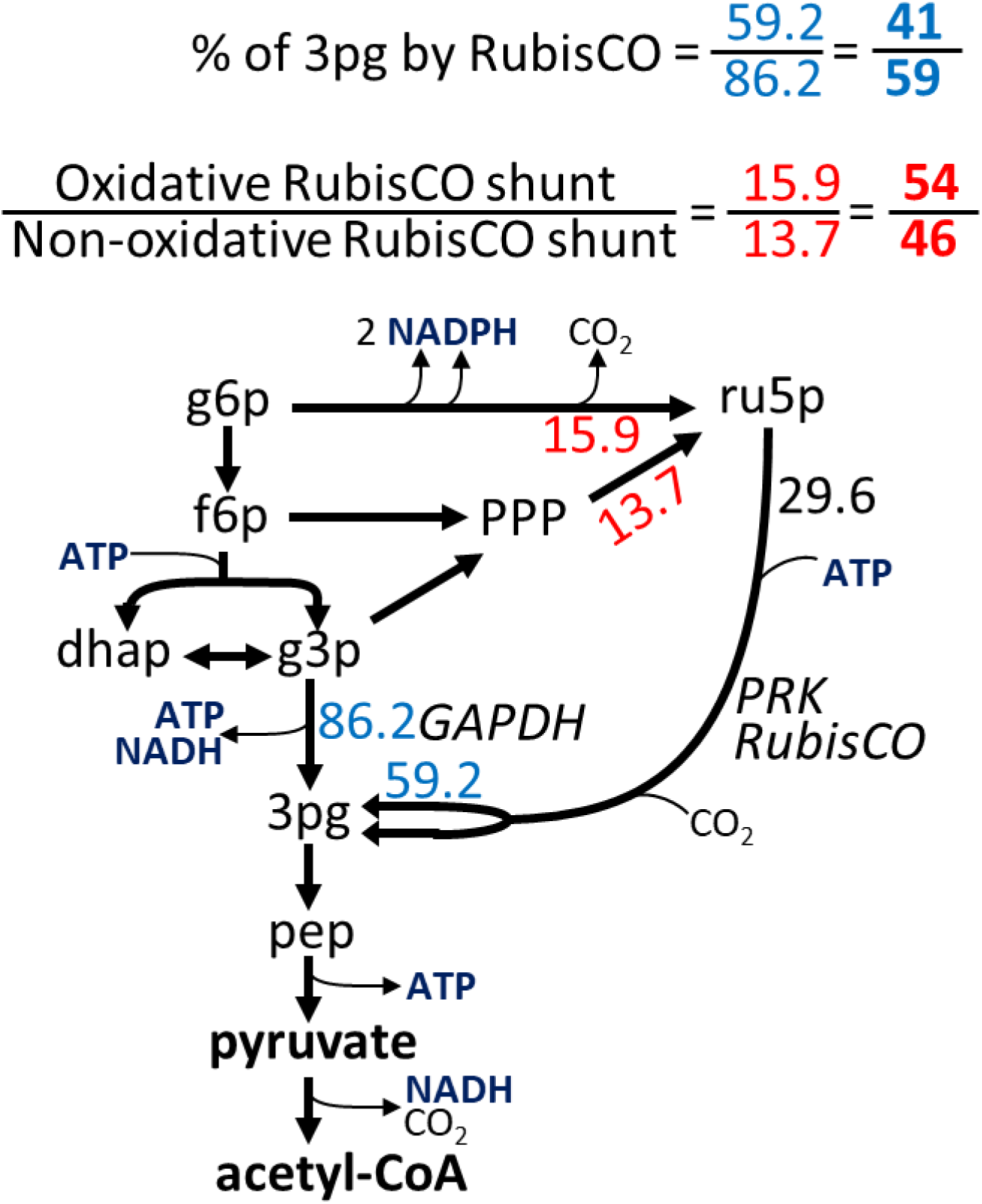
Flux ratios that describe the contribution of RubisCO to seed metabolism. Shown is the conversion of glucose 6-phosphate (g6p) to acetyl-CoA via glycolysis and the bypass variants of the RubisCO shunt. Select FBA flux values are reproduced from Table 4 and the proportions in which 3-phosphoglycerate (3pg) is formed by glycolysis or RubisCO are highlighted in blue. The oxidative and non-oxidative RubisCO shunt differ in that ribulose-5-phosphate (ru5p) is formed by different pathways (red numbers), while the section between ru5p and 3pg is common.

Fig. 2D shows an additional type of glycolytic shunt that is obtained if npGAPDH is active (pathway 4 in Table 2). Like in the oxidative RubisCO shunt, npGAPDH proceeds as a bypass to the GAPDH and phosphoglycerate kinase (PGK) glycolytic steps. While in conventional glycolysis the GAPDH and PGK work together to oxidize glyceraldehyde-3-phosphate (g3p) to 3pg with the production of NADH and ATP, npGAPDH forms 3pg and NADPH without generating ATP. npGAPDH is present and conserved among higher plants (Piattoni *et al*., 2011; Piattoni *et al*., 2013). Although npGAPDH is known to be localized to the cytosol (Piattoni *et al*., 2011; Piattoni *et al*., 2013), it might still be relevant to contribute NADPH to plastidic FAS if the produced NADPH can be transferred into the plastid, e.g. via the malate-oxaloacetate shuttle mechanism (Selinski and Scheibe, 2019). Indeed, presence of such shuttle mechanisms in *Bna572+* explains why the model reduction process identified npGAPDH as active in glucose-to-TAG pathway schemes.

### The effect of glycolytic shunts can be described as an ATP driven transhydrogenation

Overall, pathways 2 to 4 exhibit the same overall conversion efficiency of 65.5% (Table 2) and the respective glycolytic shunts are functional in correcting NADH overproduction at the glycolytic NADH-GAPDH step by replacing it in some way with NADPH. This effect is further illustrated in Table 3 by comparing the overall stoichiometries of the malate shunt, the oxidative RubisCO shunt, and the npGAPDH shunt with those of the respective bypassed glycolytic segments. For example, considering the conversion of phosphoenolpyruvate (pep) to pyruvate (pyr), the summary stoichiometry for the malate shunt is compared to the stoichiometry of pyruvate kinase (Table 3). The difference as a stoichiometry is equal to a transhydrogenation of NADH to NADP at the expense of hydrolyzing one mole ATP to ADP and inorganic phosphate (Table 3). This means that, relative to the bypassed glycolysis section, transhydrogenation is achieved at the cost of ATP. The same result is obtained when the same analysis is performed for the oxidative RubisCO shunt (glucose 6-phosphate to 3-phosphoglycerate) and for the npGAPDH shunt (glyceraldehyde-3-phosphate to 3-phospho-glycerate) (Table 3).

**Table 3:** Summary stoichiometries of glycolysis shunts in the small-scale model. Summary stoichiometries of glycolysis sections and respective shunts were derived and then, in each case, the shunt stoichiometry was subtracted from the stoichiometry of the respective glycolysis section (Supplemental Table S6). Relative to glycolysis, the summary stoichiometries of the shunts cause net transhydrogenation of NADH to NADP at an energetic cost of 1 ATP.

| pathway | Combined stoichiometries | Number of participating reactions |
| --- | --- | --- |
| **Glycolysis section: phosphoenolpyruvate (pep) → pyruvate (pyr)** |  |  |
| Glycolysis (1) | $1 \text{ pep} + 1 \text{ adp} + 1 \text{ h} \rightarrow 1 \text{ pyr} + 1 \text{ atp}$ | 1 |
| Malate shunt (2) | $1 \text{ pep} + 1 \text{ h}_2\text{o} + 1 \text{ nadh} + 1 \text{ nadp} \rightarrow 1 \text{ pyr} + 1 \text{ nad} + 1 \text{ nadph} + 1 \text{ pi}$ | 3 |
| (2) – (1) | $1 \text{ nadp} + 1 \text{ nadh} + 1 \text{ atp} + 1 \text{ h}_2\text{o} \rightarrow 1 \text{ nadph} + 1 \text{ nad} + 1 \text{ adp} + 1 \text{ pi} + 1 \text{ h}$ | |
| **Glycolysis section: glucose 6-phosphate (g6p) → 2 x 3-phospho-D-glycerate (3pg)** |  |  |
| Glycolysis (1) | $1 \text{ g6p} + 2 \text{ nad} + 1 \text{ adp} + 2 \text{ pi} \rightarrow 2 \text{ 3pg} + 2 \text{ nadh} + 1 \text{ atp} + 3 \text{ h}$ | 6 |
| Oxidative RubisCO shunt (2) | $1 \text{ g6p} + 2 \text{ nadp} + 2 \text{ h}_2\text{o} + 1 \text{ atp} \rightarrow 2 \text{ 3pg} + 2 \text{ nadph} + 1 \text{ adp} + 5 \text{ h}$ | 5 |
| (2) – (1) | $2 \text{ nadp} + 2 \text{ nadh} + 2 \text{ atp} + 2 \text{ h}_2\text{o} \rightarrow 2 \text{ nadph} + 2 \text{ nad} + 2 \text{ adp} + 2 \text{ pi} + 2 \text{ h}$ | |
| **Glycolysis section: D-glyceraldehyde-3-phosphate (g3p) → 3-phospho-D-glycerate (3pg)** |  |  |
| Glycolysis (1) | $1 \text{ g3p} + 1 \text{ nad} + 1 \text{ pi} + 1 \text{ adp} \rightarrow 1 \text{ h} + 1 \text{ nadh} + 1 \text{ atp} + 1 \text{ 3pg}$ | 2 |
| npGAPDH* (2) | $1 \text{ g3p} + 1 \text{ h}_2\text{o} + 1 \text{ nadp} \rightarrow 2 \text{ h} + 1 \text{ nadph} + 1 \text{ 3pg}$ | 1 |
| (2) – (1) | $1 \text{ nadp} + 1 \text{ nadh} + 1 \text{ atp} + 1 \text{ h}_2\text{o} \rightarrow 1 \text{ nadph} + 1 \text{ nad} + 1 \text{ adp} + 1 \text{ pi} + 1 \text{ h}$ | |

Abbreviations: npGAPDH, non-phosphorylating glyceraldehyde-3-phosphate dehydrogenase; g3p, D-glyceraldehyde-3-phosphate; g6p, D-glucose-6-phosphate; pep, phosphoenolpyruvate; pi, phosphate; pyr, pyruvate; x3pg, 3-phospho-D-glycerate
\* Non-phosphorylating glyceraldehyde-3-phosphate dehydrogenase ALDH11A3 (At2g24270)

**Table 4:** Comparison of ^13^C-MFA flux results and FBA flux prediction. Characteristic fluxes are reproduced in flux units relative to the total uptake of sucrose in hexose units (100). ^13^C-MFA flux results are taken from a previous study where *B. napus* 3170 embryos were *in-vitro* cultured (Hay *et al*., 2014). For FBA prediction using *Bna572+*, light flux, ATP and NADPH maintenance fluxes were adjusted so that the FBA model matches the CCE and relative OPPP flux from the ^13^C-MFA study (See “Methods”). Then flux was predicted by a three-level optimization by 1) minimizing the total carbon uptake relative to a fixed biomass flux, 2) maximizing glucose-6-phosphate dehydrogenase and 3) predicting optimum flux range for reactions (Flux variability).

| Reaction (objective function, reactions in *Bna572+*) | $^{13}\text{C}$-MFA | FBA predicted flux range |
| --- | --- | --- |
| Total hexose uptake ($\text{SUT}_{\text{ac}}$) | 100 | 100 |
| Ala uptake ($\text{Ala\_H\_sym}_{\text{ac}}$) | $7.57 \pm 1.30$ | 13.19 |
| Gln uptake ($\text{Gln\_H\_sym}_{\text{ac}}$) | $6.66 \pm 0.23$ | 4.17 |
| Carbon Conversion Efficiency | $81.09 \pm 0.68$ | 81.13 |
| Glucose 6-phosphate / phosphate antiporter ($\text{G6P\_c\_Pi\_p\_anti}$) | $75.10 \pm 38.97$ | [-9.53, 226.16] |
| phosphoenolpyruvate/ phosphate antiporter ($\text{PEP\_c\_Pi\_p\_anti}$) | $5.59 \pm 22.13$ | [-424.75, 287.27] |
| glucose-6-phosphate dehydrogenase ($\text{G6PDH}_{\text{c}} + \text{G6PDH}_{\text{p}}$) | $15.92 \pm 4.09$ | 15.92 |
| ribulose-bisphosphate carboxylase oxygenase ($\text{RuBisC}_{\text{p}}$) | $32.41 \pm 3.93$ | 29.59 |
| ribulose-5-phosphate from non-oxidative section of pentose phosphate pathway ($-\text{vPPepi\_p} - \text{vPPiso\_p} - \text{vPPepi\_c}$) | $16.49 \pm 4.78$ | 13.67 |
| Glyceraldehyde 3-phosphate dehydrogenase ($\text{NADGAPDH}_{\text{p}} + \text{NADGAPDH}_{\text{c}} + \text{ALDH}_{\text{c}} - \text{NADPGAPDH}_{\text{p}}$) | $81.18 \pm 7.77$ | 86.19 |
| phosphoenolpyruvate carboxylase ($\text{PEPC}_{\text{c}}$) | $6.33 \pm 0.34$ | 6.96 |
| NADPH-malic enzyme ($\text{ME}_{\text{c}}$) | $0.38 \pm 0.28$ | 0.00 |
| NADPH-malic enzyme ($\text{ME}_{\text{p}}$) | $0.06 \pm 0.08$ | 0.00 |
| pyruvate transport cytosol – plastid ($\text{H\_PYR\_sym}_{\text{cp}}$) | $17.72 \pm 4.07$ | [0.00, 133.87] |
| Pyruvate dehydrogenase complex – plastid ($\text{PDH}_{\text{p}}$) | $129.25 \pm 1.40$ | 128.67 |
| pyruvate transport cytosol – mitochondria ($\text{PTP}_{\text{cm}}$) | $8.25 \pm 0.74$ | 10.73 |
| citrate synthase ($\text{CS}_{\text{m}}$) | $9.90 \pm 0.24$ | 10.73 |
| isocitrate dehydrogenase ($\text{IDH}_{\text{m}} + \text{IDH}_{\text{c}} + \text{IDH}_{\text{p}}$) | $-1.13 \pm 0.13$ | 0.10 |
| oxoglutarate dehydrogenase ($\text{OGDH}_{\text{m}}$) | $1.41 \pm 0.20$ | 0.00 |

The ATP expenditure for transhydrogenation is probably necessary due to reaction thermodynamics. The two redox cofactors NAD(H) and NADP(H) usually are assumed to fulfill different roles in metabolism (Smith *et al*., 2021; VanLinden *et al*., 2015). In plant tissues, as in other organisms, NADH is thought to primarily serve oxidative phosphorylation in the mitochondria, leading to ATP formation. In contrast, NADPH is thought to be a reducing agent for anabolic pathways (Smith *et al*., 2021; VanLinden *et al*., 2015). Consistent with this division of labor, the NADPH/NADP redox ratio is typically significantly higher than the NADH/NAD ratio (Harold, 1986; Heineke *et al*., 1991; Igamberdiev and Gardeström, 2003). If the NADPH/NADP redox couple is more reduced, the transfer of reducing equivalents from NADH to NADP should be energetically unfavorable unless driven by an additional energy expenditure.

The network topography of the small-scale model doesn’t allow NADH-to NADP transhydrogenation only at the cost of 1 ATP.

### Mitochondrial oxidative phosphorylation contributes significantly to the cofactor budget

As mentioned above, in the unbalanced scenario in Fig. 2A, a significant activity of oxidative phosphorylation discharges NADH overproduction. The balanced scenarios (Fig. 2B-D) rely on this mitochondrial process as well, but here to meet the biosynthetic ATP demand. In all three balanced scenarios 3.67 moles of the generated NADH is converted into 7.33 ATP by oxidative phosphorylation (Fig. 2B-D), which amounts to 44% and 73.3% of the biosynthetic NADH and ATP demands, respectively. The stoichiometries of Fig. 2 are based on a P/O ratio of 2, which has been assumed to be a realistic value for the oxidation of cytosolic NADH (Lambers, 1997). Since exact P/O ratios in mitochondrial oxidative phosphorylation might vary with conditions or are matter of debate (Ferguson, 1986; Hinkle, 2005), the glucose to TAG conversion was re-calculated for a range of P/O ratios between 1 and 4 (Table S7, Fig. S4). According to this wide range, the mitochondrial NADH consumption and ATP production amount to between 20.9 and 39.8% of the NADH demand and 55.0 and 88.0% of the ATP demand, respectively (Fig. S4). Thus, in a purely heterotrophic context as assumed in the small-scale model, plastidic FAS might be substantially dependent on mitochondrial oxidative phosphorylation. Accordingly, the existence of shuttle mechanisms for the movement of NADH reducing equivalents from the plastid into mitochondria under certain conditions is supported by literature. Mutational analysis in Arabidopsis has shown that PLASTIDIAL NAD-DEPENDENT MALATE DEHYDROGENASE (plNAD-MDH), the chloroplastic DICARBOXYLATE TRANSPORTER 1 (DiT1) and MITOCHONDRIAL MALATE DEHYDROGENASE 1 (mMDH1) are components of a malate/oxaloacetate shuttle mechanism that can move surplus NADH reducing equivalents from the chloroplast into the mitochondria when fatty acid synthesis is partially blocked (Zhao *et al*., 2018; Zhao *et al*., 2020). The mechanism seems to be relevant during seed development since a knockout of plNAD-MDH is embryo lethal (Selinski *et al*., 2014). In addition, biochemical evidence suggests that, particularly in heterotrophic organs, plastidic FAS is to a large extent dependent upon provision of ATP from the cytosol (Neuhaus and Emes, 2000; Rawsthorne, 2002). Furthermore, in Arabidopsis, double knockout of the two isoforms of the ATP/ADP transporter of the plastidic inner envelope (NTT1, NTT2) resulted in a significant reduction in seed oil, and it was concluded that ATP import into the plastids exerts significant control over lipid synthesis during Arabidopsis seed development (Reiser *et al*., 2004). Similar knockout and overexpression studies also suggested that the NTT is important for seed oil accumulation in *Brassica napus* as well (Hong *et al*., 2022; Xia *et al*., 2022). Altogether, the dependence of TAG synthesis on mitochondrial activity deduced from the small-scale model has support from literature.

### Assessing glycolytic shunts in a wider context of seed metabolism

Having demonstrated with a small-scale model that glycolysis-based conversion of glucose to triacylglycerol (TAG) leads to an intrinsically unbalanced cofactor profile—and that a balanced conversion can be achieved only when one or more of the three described glycolytic shunts are active—it becomes essential to assess whether these findings hold true within the broader framework of cellular metabolism in developing oilseeds. In the larger metabolic context of seed development, TAG biosynthesis represents only one of several key processes. Other major biosynthetic demands include the production of proteins, cell wall polysaccharides, starch, and nucleic acids. Furthermore, in a realistic FBA model, a significant portion of the cell’s ATP and NADPH budget must be allocated to maintenance processes that do not directly contribute to biomass accumulation or the synthesis of storage compounds (Cheung *et al*., 2013; De Vries, 1974; Thiele and Palsson, 2010). Therefore, any surplus ATP generated during the conversion of sugars to TAG could potentially support the other anabolic activities or being consumed by the ATP maintenance demands. From this the question arises whether the glycolytic shunts are predicted to be relevant in the larger metabolic context of a FBA model for seed metabolism.

To explore TAG synthesis in a realistic physiological context, the *Bna572+* model was adjusted using experimentally derived parameters for *Brassica napus in-vitro* cultured embryos and results from ^13^C-MFA (Hay *et al*., 2014). Specifically, the contributions of the photosynthetic ATP and NADPH generation as well as ATP and NADPH maintenance demands to the energy budget in the model were adjusted so that the overall CCE and the relative OPPP flux of the ^13^C-MFA flux results were matched (see “Methods” for details). After these adjustments, when using a two-level FBA procedure where total carbon uptakes are minimized and then the OPPP flux is maximized, key ^13^C-MFA derived fluxes, including OPPP flux, are largely in agreement FBA predicted ones (Table 4). Fluxes associated with the RubisCO shunt (*RuBisC_p_*, Table 4) and the tricarboxylic acid cycle (reactions *CS_m_*, *IDH*, *ME*, *OGDH_m_*, *PEPC_c_*, Table 4) were closely matched by the FBA prediction. The very low fluxes through isocitrate dehydrogenase (*IDH*) and oxoglutarate dehydrogenase (*OGDH*) (Table 4) indicate that the tricarboxylic acid cycle is only used to a limited extent for catabolism/generation of energy cofactors, which is also consistent with previous flux studies in *B. napus* (Schwender *et al*., 2015; Schwender *et al*., 2006). In addition, consistent with the flux through malic enzyme isoforms being very low in the ^13^C-MFA study, the FBA simulation procedure predicts the cytosolic and plastidic isoforms of malic enzyme to be inactive (Table 4).

In the FBA scenario of Table 4, the RubisCO shunt is the preeminent glycolytic shunt. As in previous studies, the relative activity of the RubisCO shunt can be well characterized by a flux ratio around 3pg, shown in blue numbers in Fig. 3 (Schwender *et al*., 2004). Fig. 3 shows that 41% of the 3pg processed toward FAS is generated via RubisCO, which comes close to the previously determined range of 37–51% for embryos cultured under the same conditions (Schwender *et al*., 2004). In the FBA simulated photoheterotrophic environment, both the oxidative and the non-oxidative RubisCO shunt are involved in almost equal proportions (Fig. 3, red numbers). Both variants bypass NADH generation at the glycolytic GAPDH step, but only the non-oxidative RubisCO shunt has the advantage of improving the carbon efficiency of the overall conversion by re-fixing CO2 released at the PDH step (Fig. 3).

### In oilseeds glycolytic shunts are predicted to be active above about 30% TAG

With the adjustment of *Bna572+* to experimental findings of *in-vitro* cultured *B. napus* embryos, further predictions about the role of the glycolytic shunts in an oilseed can be made. If the shunts are required for TAG synthesis in the context of the larger metabolic context with multiple seed storage products, their usage should be predicted to increase with seed oil content. To test this, biomass composition was simulated for a continuously increasing TAG content, with the proportion of all other seed biomass compounds proportionally reduced, while the adjustments to biomass flux rate, light uptake flux, and ATP and NADPH maintenance requirements remained unchanged (Fig. 4). As a result, with increasing TAG content, the overall conversion of organic medium substrates to seed biomass becomes less carbon efficient (Fig. 4), which is expected due to the higher energy density of TAG compared to other biomass compounds. When the trade-off simulation was repeated each time by maximizing the oxidative RubisCO shunt, the npGAPDH shunt, or the malate shunt in the second-level optimization, it was found that the three can work alternatively: It is predicted that all three glycolytic shunts can be active above 30% TAG and from there increase monotonically and to a similar extent (Fig. 4). The explanation for this increase is that with increasing TAG content, the conversion of sucrose to TAG becomes an increasingly dominant process, whereby the above-described oversupply of NADH by glycolysis becomes quantitatively more pronounced. In Table 3, it was shown that the activity of glycolytic shunts to compensate for excess NADH causes an ATP expenditure compared to the pure glycolytic conversion. Using the FBA simulation framework, model sensitivity analysis confirms that the cost for NADH to NADP transhydrogenation is 1 ATP. This applies particularly for the range of TAG content above 30% where the glycolytic shunts become active (Fig. 4). Although a TAG content of 90% is likely unattainable, the trade-off simulation in Fig. 4 was extended to this value (Fig. 4). This is to show that when TAG is the only product, the relative contributions of the glycolytic shunts are comparable to those observed with the small-scale model (Fig. 2). In the small-scale model, 66% of the glycolytic formation of pyruvate (Fig. 2B) and 3-phosphoglycerate (3pg, Fig. 2C, D) are bypassed. In similar, in Fig. 4, at a TAG level of 90%, the oxidative RubisCO shunt and the np-GAPDH shunt bypasses 66.4%, while the malate shunt bypasses 65.3% of glycolytic flux. Finally, if the operation of the glycolytic shunts is required to rectify a cofactor imbalance in the sugar-to-TAG conversion, then one should expect that knockout of all three shunts cause a less efficient conversion and an ATP surplus. Consistently, if we block the glycolytic shunts (ME, npGAPDH and RuBisCO) in Bna572+, repeated simulation along the TAG range from 10 to 90 % reveals a less efficient conversion (reduced CCE) and an ATP surplus that rises with increasing TAG levels (Fig. 4). At 90 % TAG, close to 70 % of the total of ATP generated in the network is dissipated by metabolic processes that hydrolyze ATP without any use.

**Fig. 4.**
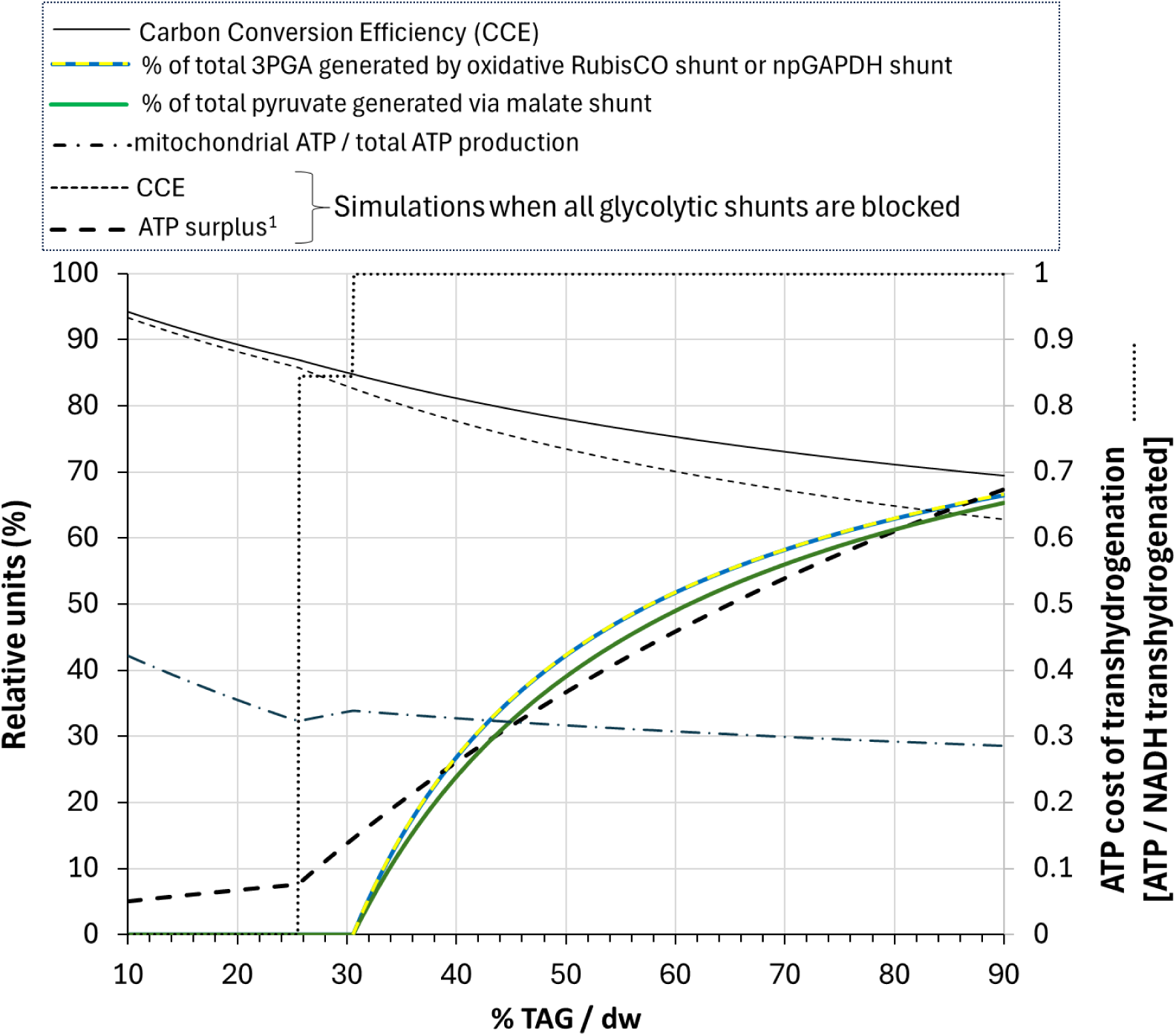
Dependence of the predicted activity of glycolytic shunts on TAG content. The predictions are based on the adjustments of light, ATP and NADPH maintenance fluxes in *Bna572+* to match experimental ^13^C-MFA results at 40% TAG (w/dw) (Table 4). Then FBA flux predictions were repeated for a tradeoff between TAG and the other biomass components. Footnotes: ^1^, ATP surplus as percentage of total ATP production when all the glycolytic shunts are blocked by inactivating reactions ME_c_, ME_p_, ALDH_c_ and RuBisC_p_.

In summary, the modeling analysis presented here integrates prior experimental observations regarding the operation of unconventional glycolytic shunts in oilseeds during lipid biosynthesis. It provides a theoretical framework for understanding and predicting the functionality of these pathways in developing seeds. Among these shunts modeled here, there is clear experimental evidence supporting the operation of both the RubisCO shunt and the malate shunt during seed development (O’Grady *et al*., 2012; Sagun *et al*., 2023). In contrast, the non-phosphorylating GAPDH (np-GAPDH) shunt remains more hypothetical at this stage. The pathway analysis principles employed in this study also suggest the potential existence of additional glycolytic bypasses, such as the phosphoketolase shunt, which has been effectively engineered in oleaginous yeasts (Qiao *et al*., 2017).

This modeling framework highlights the need for further experimental investigations, particularly concerning cofactor imbalances during oil biosynthesis. Such studies are inherently challenging due to the technical difficulty of analyzing redox metabolism and tracking reducing equivalents *in vivo* (Smith *et al*., 2021). However, promising approaches include the use of fluorescent protein-based biosensors to monitor subcellular NAD(P)H/NAD(P)+ dynamics in plants (Smith *et al*., 2021).

With respect to the RubisCO shunt, recent findings suggest its role in developing Arabidopsis seeds may go beyond simply enhancing carbon-use efficiency via CO₂ refixation. Specifically, disruption of the RubisCO shunt through loss of phosphoribulokinase (PRK) has been shown to severely impair lipid accumulation (Deslandes-Herold *et al*., 2023). Because complete PRK knockout is lethal to plants, a genetic complementation strategy was employed: PRK expression was restored using a fluorescently labeled construct, allowing homozygous *prk*/*prk* embryos to be isolated from developing siliques of heterozygous PRK/*prk* plants. Mature *prk*/*prk* embryos exhibited a 34% reduction in total fatty acid content compared to their PRK-expressing siblings, a significantly greater impact than would be predicted from theoretical estimates of the shunt’s contribution. Additional support for a broader role of the RubisCO shunt comes from metabolic phenotypes observed in *prk*/*prk* embryos, including increased starch accumulation late in development and elevated levels of hexose phosphate pools. These findings hint at a disruption of glycolytic flux and carbon reallocation toward starch. One plausible underlying cause is a redox cofactor imbalance (Deslandes-Herold *et al*., 2023), as proposed in this study. This hypothesis could be further tested by assessing oxidative stress markers, such as reactive oxygen species, as indirect indicators of redox perturbation.

This work also raises the intriguing possibility that the RubisCO shunt operates in heterotrophic contexts. So far, its function has been framed within a photoheterotrophic setting, where its contribution to CO₂ refixation enhances carbon efficiency in tandem with ATP and NADPH from photosynthetic light reactions (Schwender *et al*., 2004). However, there is evidence that the RubisCO shunt may also function in non-green, heterotrophic seeds. For example, despite lacking chlorophyll, castor bean seeds (*Ricinus communis*) exhibit significant RubisCO activity. Comparative enzyme assays indicate that RubisCO capacity in developing endosperm is sufficient to sustain fatty acid biosynthetic fluxes (Simcox *et al*., 1977). Similarly, in non-green seeds of sesame (*Sesamum indicum*), RubisCO small subunit transcripts are expressed at levels comparable to those in green seeds of Arabidopsis during seed development (Suh *et al*., 2003). In *Prunus sibirica*, notable expression of PRK and RubisCO subunits has also been observed in its non-green oil-accumulating seeds (Wang *et al*., 2019). Although *Brassica napus* seeds possess photosynthetic activity and RubisCO expression (Schwender *et al*., 2004), constraint-based modeling suggests that inner cotyledon tissue during seed development grows under largely heterotrophic conditions (Borisjuk *et al*., 2013). Thus, a heterotrophic role for the non-oxidative RubisCO shunt in green seeds such as *B. napus* remains both plausible and relevant.

## Supplementary data

Fig. S1. Identification of pathway schemes that allow transhydrogenation from NADH to NADP at no cost in Bna572+

Fig. S2. Model adjustment to experimental flux values based on a pareto front for a tradeoff between ATP and NADPH maintenance cost.

Fig. S3. Model adjustment to experimental flux values for different levels of light flux.

Fig. S4. Dependence of balanced glucose-to-TAG conversions on the P/O ratio.

Table S1: Dependence of OPPP flux in FBA simulations on irreversibility constraints in Bna572+.

Table S2. Reduced model version of Bna572+.

Table S3. Set of reactions included in the small-scale model for the conversion of glucose to triacylglycerol.

Table S4. Metabolites included in the minimal model for the conversion of glucose to triacylglycerol.

Table S5. Extreme pathway analysis results for the small-scale model.

Table S6. Summary stoichiometries of glycolysis bypasses in the small-scale model.

Table S7. Dependence of characteristics of the small model from the P/O ratio.

Supplemental Material 1: Chemical mass- and redox balance for the conversion of glucose into reduced compounds.

Supplementary File 2: Model and code files.

## Author contributions

JS conceived the project, performed the research and wrote the article.

## Data availability

The original contributions presented in the study are included in the article/Supplementary Material, further inquiries can be directed to the corresponding author.

## Abbreviations

CCE: carbon conversion efficiency
COBRA: Constraint-Based Reconstruction and Analysis
FA: fatty acid
FAS: fatty acid synthesis
IDH: isocitrate dehydrogenase
OPPP: oxidative pentose-phosphate pathway
PEP: phosphoenolpyruvate
PGA: phosphoglycerate
RubisCO: Ribulose-1,5-bisphosphate carboxylase/oxygenase
TCA: tricarboxylic acid
TH: transhydrogenation.

